# Cross-Domain Text Mining of Pathophysiological Processes Associated with Diabetic Kidney Disease

**DOI:** 10.1101/2024.01.10.575096

**Authors:** Krutika Patidar, Jennifer H. Deng, Cassie S. Mitchell, Ashlee N. Ford Versypt

## Abstract

Diabetic kidney disease (DKD) remains a significant burden on the healthcare system and is the leading cause of end-stage renal disease worldwide. The pathophysiology of DKD is multifactorial and characterized by various early signs of metabolic impairment, inflammatory biomarkers, and complex pathways that lead to progressive kidney damage. New treatment prospects rely on a comprehensive understanding of disease pathology. The study aimed to identify signaling drivers and pathways that modulate glomerular endothelial dysfunction in DKD via cross-domain text mining with SemNet 2.0. The open-source literature-based discovery approach, SemNet 2.0, leverages the power of text mining 33+ million PubMed articles to provide integrative insight into multiscalar and multifactorial pathophysiology. A set of identified relevant genes and proteins that regulate different pathological events associated with DKD were analyzed and ranked using normalized mean HeteSim scores. High-ranking genes and proteins intersecting three domains—DKD, immune response, and glomerular endothelial cells—were analyzed. The top 10% of ranked concepts mapped to the following biological functions: angiotensin, apoptosis, cell-cell function, cell adhesion, chemotaxis, growth factor signaling, vascular permeability, nitric oxide response, oxidative stress, cytokine response, macrophage signaling, NFκB factor activity, TLR signaling, glucose metabolism, inflammatory response, ERK/MAPK signaling, JAK/STAT signaling, T-cell mediated response, WNT signaling, renin angiotensin system, and NADPH response. High-ranking genes and proteins were used to generate a protein-protein interaction network. This comprehensive analysis identified testable hypotheses for interactions or molecules involved with dysregulated signaling in DKD, which can be further studied through biochemical network models.

## 1 Introduction

Diabetic kidney disease (DKD) is a major microvascular complication in the kidney that affects patients with type I diabetes and type II diabetes. DKD can lead to a decline in kidney function and has the potential to develop into chronic kidney disease or end-stage renal disease (ESRD) [1]. A high clinical and socio-economic impact of DKD is burdensome because of the risk of progression to ESRD and other related comorbidities [2]. The pathophysiology of DKD is multifactorial and characterized by a critical metabolic impairment, uncontrolled inflammatory response, increased apoptosis, and tissue fibrosis [2]. In diabetes, aberrant glucose metabolism leads to dysregulated immune response and signaling [3]. The dysregulated signaling response leads to progressive kidney damage through loss of glomerular endothelial fenestrations, thickening of the basement membrane, detachment of podocyte foot processes, mesangial matrix expansion, and glomerular fibrosis [3–5].

Various mathematical models have demonstrated the renal physiological and pathophysiological processes involved in DKD. Hyperglycemia-induced podocyte injury in DKD has been previously modeled using the local renin-angiotensin system in renal podocyte cells [6]. Some mechanistic evidence is included in a mathematical model about the disease etiology of glomerular fibrosis in DKD [5]. We have also previously modeled a protein-protein interaction network (Figure 1) between pro-inflammatory macrophages and glomerular endothelial cells (GEC) in the kidney using logic-based ordinary differential equations to understand the effect of inflammation and immune response on the progression of DKD [7]. This multi-cell network was a manually curated, simplified network of pathway interactions and signaling molecules that affect glomerular endothelial fenestrations in the diabetic kidney. However, these biochemical network models only incorporated a subset of the relevant pathways, interactions, or molecules governing DKD progression.

**Figure 1.**
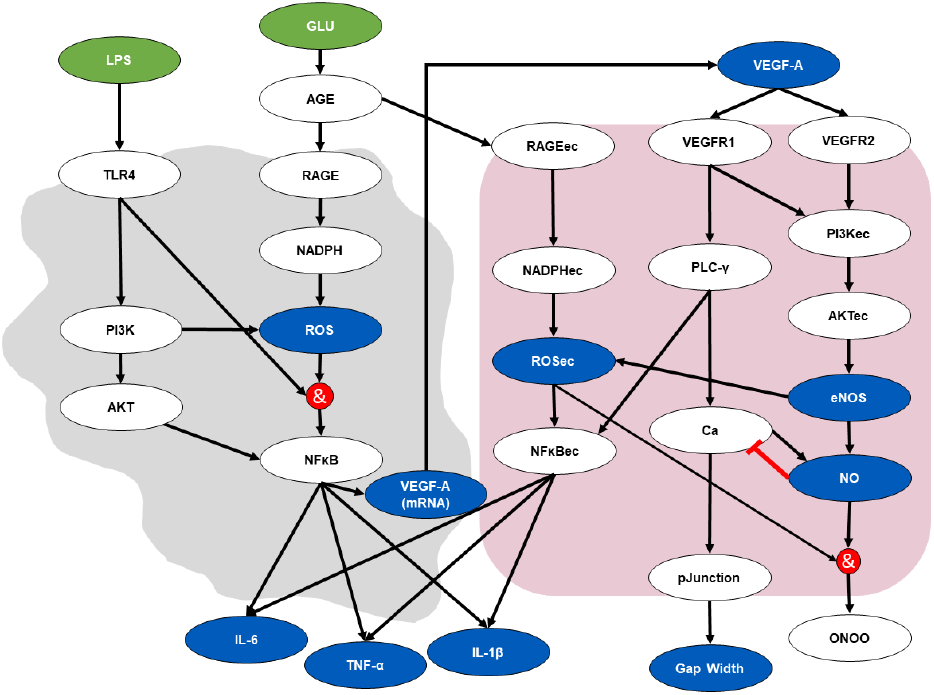
A multi-cellular protein-protein interaction network of crosstalk between macrophages (left) and glomerular endothelial cells (right) stimulated with glucose (GLU) and lipopolysaccharide (LPS), a pro-inflammatory stimulus, was created in our previous work through manual curation of the literature [7]. Green nodes (ovals) are input nodes, blue nodes are output nodes, and white nodes are regulatory nodes. Black arrows are activating interactions, a red line with a flat-head arrow is an inhibiting interaction, and red circles indicate logic *AND* gates. An *OR* logic rule connects two or more edges to a subsequent node throughout the network unless indicated otherwise by an *AND* logic gate. The subscript (ec) denotes an intracellular species expressed in endothelial cells. IL-6, TNF-α, IL-1β, and VEGF-A are protein levels expressed in extracellular space. ROS, ROSec, VEGF-A (mRNA), and NO are expressed within the cells. The Gap Width node denotes a fractional change in the glomerular endothelial cell fenestration size. The pJunction node represents the phosphorylated junction protein levels. TLR: toll-like receptor. AGE: advanced glycation end product. RAGE: receptor of advanced glycation end product. NADPH: nicotinamide adenine dinucleotide phosphate. NFκB: nuclear factor kappa B. IL: interleukin. TNF: tumor necrosis factor. PI3K: phosphoinositide 3-kinases. AKT: serine/threonine-specific protein kinases. ROS: reactive oxygen species. VEGF: vascular endothelial growth factor. VEGFR: vascular endothelial growth factor receptor. PLC: phospholipase C. NO: nitric oxide. ONOO: peroxynitrite. eNOS: endothelial nitric oxide synthase. Ca: calcium. Image reused from [7].

Determining the relevant pathway interconnections to include in a mathematical model or a protein-protein interaction network via manual literature curation is inherently challenging. A human cannot read all possible sources of important relationships. Moreover, signaling molecules in the disease pathway may be located further away from the target molecule or event of interest within the scientific literature [8]. Thus, a manual literature search may be insufficient in finding relationships not only within the DKD literature but also in cross-domain literature (cardiology, endocrinology, immunology, etc.) where relevant concepts and relationships may reside. We contend a holistic understanding of relevant dysregulated pathways and molecules associated with DKD can be deduced from literature using advanced high-throughput artificial intelligence approaches to literature-based discovery (LBD) like SemNet 2.0 [9]. LBD approaches enabled by text-mining infer disparate sources of information [10] at a scale not otherwise possible.

SemNet 2.0 is an open-source framework comprising a knowledge graph that identifies and ranks the most important concepts to a user-defined target concept(s) (e.g., keyword). The graph is comprised of relationships extracted from 33+ million PubMed articles. The nodes are concepts defined by the Unified Medical Language System. The framework uses SemMedDB as a relationship extraction system for making the graph [11]. The unsupervised learning ranking algorithm within SemNet 2.0 examines relationship patterns in the literature to rank cross-domain concepts with respect to the user-defined concept(s) [9]. SemNet 2.0 has been used for drug repurposing of COVID-19 [12] and Parkinson’s disease [13], to identify unknown disease mechanisms of resistant hypertension following COVID-19 infection [14], and to predict adverse events from chronic tyrosine kinase inhibitor therapy in chronic myeloid leukemia [8].

In this study, we use SemNet 2.0 to identify and rank critical signaling molecules associated with glucose- and inflammation-mediated development and progression of DKD. By evaluating highly ranked concepts from over 33+ million PubMed articles, concepts or phenomena missing from current mathematical network models of DKD were identified and ranked. The artificial intelligence-assisted LBD approach provided a less biased and more comprehensive manner for integrating cross-domain knowledge into the mechanisms of DKD.

## 2 Methods

The present study used advanced text-mining techniques to identify relevant signaling molecules and their relation to glucose-mediated inflammation in DKD. Specifically, we used SemNet 2.0 to find integrated text-mined nodes across previously studied domains such as diabetes, kidney disease, immune response and inflammation, and glomerular endothelial cells. SemNet 2.0 performed a cross-domain analysis that focused on five disease domains. A pairwise search was performed across the following five domains: diabetes (DB), kidney disease (KD), immune response (IR), glomerular endothelial cell (GEC), and DKD (Figure 2). The list of user-specified target nodes of interest that are most relevant to characterize each domain can be found in Table S1. These target nodes of interest were chosen from observed interactions in the network model and published studies [7]. For instance, the binding of toll-like receptors (TLRs) is one of the key determinants of the immune response. Therefore, it was considered one of our investigation’s target nodes.

**Figure 2.**
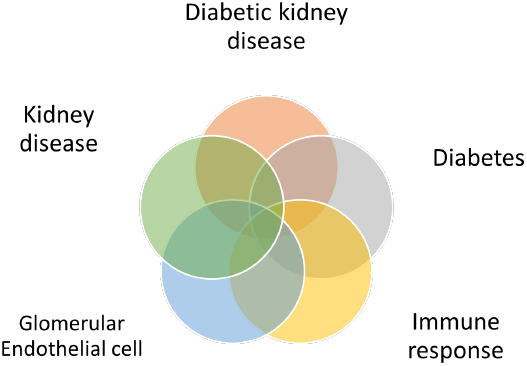
Visualization of cross-domain analysis between 5 domains: diabetes, diabetic kidney disease), immune response, glomerular endothelial cell, and kidney disease.

**Figure 3.**
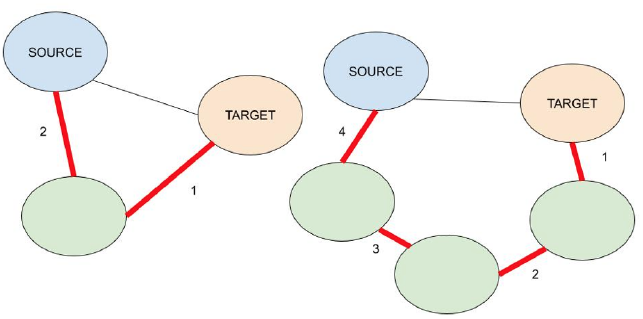
In the subgraph on the left, the source node is at a depth of 1 away, but the HeteSim metapath length is 2. On the right, the same source node is also at a depth of 1 away, but the HeteSim metapath length is 4. Increasing both metrics can drastically expand the scope of the search.

### 2.1 SemNet 2.0

SemNet 2.0 [9] is an open-source software that ingests publicly available text relationships from PubMed and the National Library of Medicine (NLM) [11] to perform LBD tasks. SemNet uses a heterogeneous semantic network to provide a consistent and valuable categorization of all concepts represented in the Unified Medical Language System (UMLS) metathesaurus, which provides a universal ontology to relate concepts from the biomedical literature [15]. SemNet 2.0 queries a biomedical knowledge graph composed of semantic triples extracted from PubMed’s 33 million abstracts. The original SemNet was proposed in 2019 by Mitchell and Sedler [16] and was later optimized by Kirkpatrick and colleagues in 2022 [9]. Each semantic triple consists of a head, relation, and tail, where the head and tail entities are the nodes, and the relation is a directed edge.

SemNet 2.0 is available as Python code and uses natural language processing to identify source nodes relevant to user-specified target nodes. The source nodes are the set of nodes that the target nodes share in common; that is, they are reachable within the search depth and metapath length, which are search parameters defined in the next section. Each node is a biomedical concept, as defined in the UMLS, with a type such as Disease or Syndrome (DSYN), Amino Acid Peptide Protein (AAPP), etc. There are 133 types and 54 relations. Each directed edge encodes a relation, such as treats, affects, inhibits, etc. SemNet calculates a metric called HeteSim to quantify the relevance between a source node and target node [9].

#### 2.1.1 Search Parameters

The user defines four inputs: the target nodes, source node types, search depth, and metapath length. Target nodes are the nodes of interest, and SemNet queries surrounding nodes that are connected to those nodes. The source node types can be restricted to certain semantic types, such as DSYN or AAPP. Search depth is the number of hops away from the source node. For a given target node, a search depth of 1 finds all adjacent nodes directly connected to it. It is ideal to increase search depth to find novel results, as connections of neighbors to target nodes are more prominent and commonly acknowledged in scientific literature. Metapath length is the total path from a target to a source node. Multiple paths can be consolidated into a single metapath based on the types of source nodes. Hypothetically, an infinite number of paths can be used to connect a target to a source node, and the metapath length can add a constraint for identifying relevant or innovative pathways.

#### 2.1.2 Calculating HeteSim

There are two ways to calculate HeteSim: deterministic and randomized. The deterministic HeteSim was used in all SemNet simulations in the present study to enhance accuracy at the expense of computational speed. HeteSim can be further characterized by an exact (deterministic) mean or approximate mean. The exact mean is found by aggregating HeteSim scores of multiple target nodes to the same source node. Approximate mean has a performance advantage over exact mean, especially for metapaths of greater length. For the simulations here, the exact HeteSim mean was used.

HeteSim is calculated by the cosine similarity between two probability vectors. Let *x* be defined as the path length from a given target node and source node. HeteSim takes the middle layer of nodes or the nodes at *x/*2 path length away from the target. From the target and source nodes, weights of 1 are distributed evenly across nodes. Each subsequent layer continually redistributes weights until the middle layer is reached. A left probability vector and a right probability vector are generated from either side. The cosine similarity is calculated between them, which is the HeteSim score. When combining results from multiple SemNet 2.0 simulations, the mean HeteSim scores are normalized and percentile ranked to adjust for differences in node count, path count, etc.

### 2.2 SemNet Simulations for DKD

We randomly selected two nodes from each domain in the pair (a total of four nodes) as input parameters, which were considered target nodes in SemNet. A total of 20 nodes were specified as “hub nodes” for making the hub node networks. Briefly, hub networks enable improved cross-domain analysis by functionally increasing the search depth in areas of the knowledge graph of chief interest [8, 14]. Ten pairwise simulation trials were performed, equating to 200 simulations. The search depth was 2, and the metapath length was 3. Due to computational limitations, if the number of source nodes exceeded 1000, a random sample of the source nodes was taken such that no more than 1000 nodes had their HeteSim scores calculated and ranked. If the number of source nodes exceeded 1500 or was less than 10, the random combination of target nodes was deemed unproductive, and a new simulation with a different set of target nodes was performed.

Some simulations had several hundreds of nodes within the *<*1500 range, but they were limited by a certain target node (e.g., “Disorder of mineral metabolism”). Throughout this study, each node or concept’s name and semantic type follow the UMLS ontology, and more information on semantic networks and types can be found in [15]. SemNet 2.0 predicted source nodes associated with the target nodes and ranked these source nodes by the optimized HetSim similarity metric [9]. The association of the source nodes to user-specified target nodes in each pairwise domain was measured using a mean HeteSim score, calculated by averaging the HeteSim score of recurring nodes.

An exhaustive list of predicted source nodes was obtained at the intersection of each pairwise domain analysis. The identified source nodes were categorized into semantic types such as genes, proteins, and enzymes. These source nodes appeared more than once in each pairwise domain analysis. Each unique source node’s occurrence was counted as the source node’s frequency and denoted as *count*. A mean HeteSim score for each source node was generated and used for further analysis of the source nodes. The top 10% of normalized and highly ranked source nodes were aggregated from the simulations. The top 10% of the source nodes were chosen based on the overall predicted relevance using the mean HeteSim score.

### 2.3 Code availability

We have provided the data and analysis files for this paper in a repository [17], which is available here: https://github.com/ashleefv/DKD_CaseStudy_SemNet2. The code leverages the open-source software SemNet 2.0 [9, 18], which is available in the following repository: https://github.com/pathology-dynamics/semnet-2.

## 3 Results

The distribution of source nodes in each semantic type for each pairwise domain analysis is shown in Figure 4. For instance, the distribution of source nodes at the intersection of both DB and DKD domains was found in the cross-domain analysis between DB and DKD (DB-DKD) domains. The intersecting nodes were either of the type gene (gngm) or type protein (aapp) in each pairwise domain (Figure 4). Therefore, nodes with semantic type gngm or aapp from the top 10% were chosen to assess further their biological themes and functional relevance in DKD progression. It was determined that the gngm and aapp semantic types were more relevant in understanding pathway interactions for the present applied study. The distribution of these top 10% source nodes in each paired domain is shown and characterized by their mean HeteSim score and count (Figure 5). The top 10% of source nodes obtained from the post-hoc analysis belonged mainly to the GEC, DKD, and IR domains. As seen in Figure 5, many source nodes were present at the intersection of GEC-IR, DKD-IR, and DKD-GEC domains relative to other pairwise domains. It was observed that the identified source nodes with a high mean Hetesim score were present at the intersection of the GEC and IR domain (Figure 5). This observation suggests that a synergistic signaling interaction between glomerular endothelial cells and an immune response is critical in the early stage of DKD progression.

**Figure 4.**
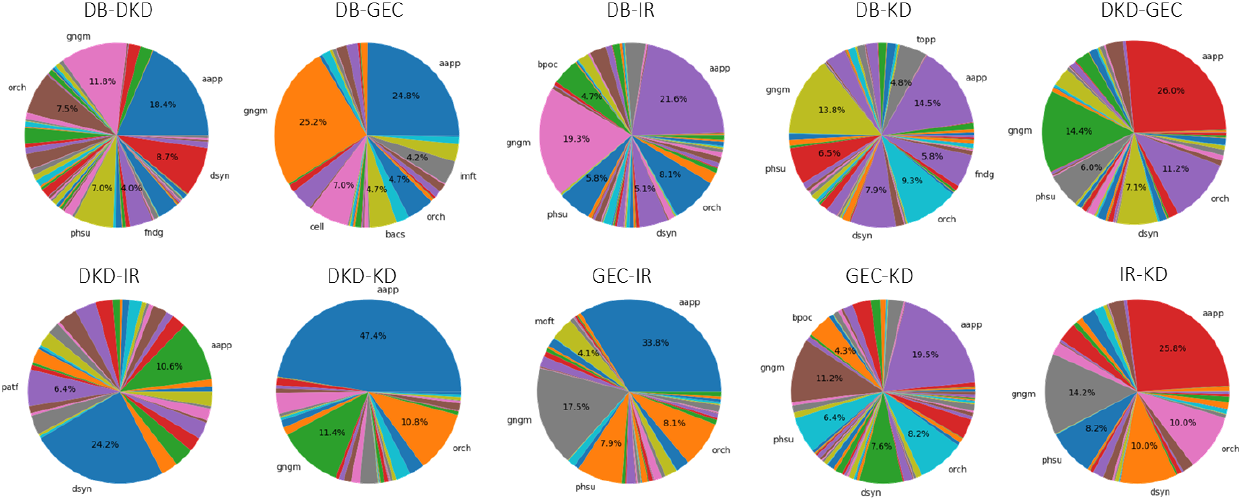
Semantic type distribution of source nodes identified at the intersection of each pairwise domain. gngm: gene or genome. aapp: amino acid, peptide, or protein. dysn: disease or syndrome. phsu: pharmacological substance. fndg: finding. imft: immunologic factor. orch: organic chemical. cell: cell. bacs: biologically active substance. patf: pathologic function. bpoc: body part, organ, or organ component. topp: therapeutic or preventive procedures. DB: diabetes domain. DKD: diabetic kidney disease domain. GEC: glomerular endothelial cell domain. IR: immune response domain. KD: kidney disease domain.

**Figure 5.**
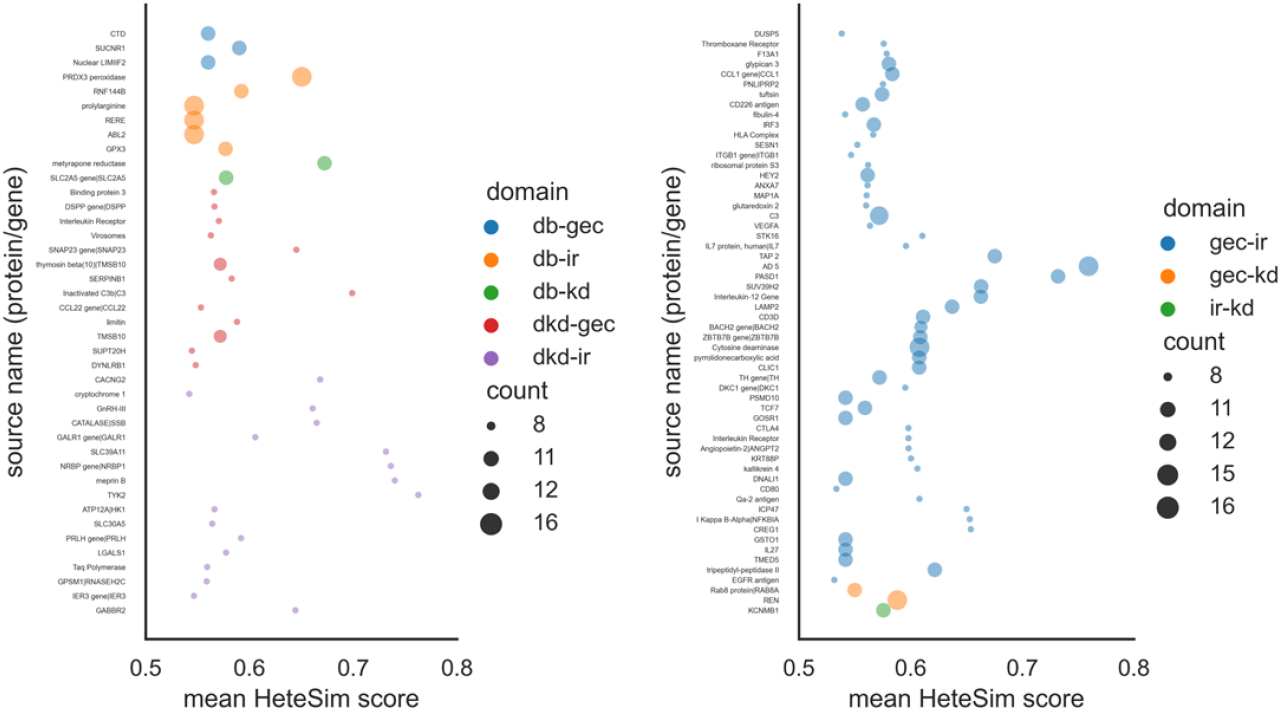
Bubble plot of source concepts identified by SemNet at the intersection of each pairwise domain. Source concepts (genes or proteins) are shown on the vertical axis, each pairwise domain is shown by bubble color, and frequency (*count* ) of each source concept is distinguished by the bubble size. For clarity, the source concepts are presented in two plots (left and right). DB: diabetes domain. DKD: diabetic kidney disease domain. GEC: glomerular endothelial cell domain. IR: immune response domain. KD: kidney disease domain. source name: protein or gene name.

The biological process or function of these genes or proteins is not available in the simulated results. Due to the vast simulation data, understanding the biological role of these source genes or proteins through a literature survey may not be feasible. Our analyses generated vast data, and thus, we searched for data-mining techniques to map the concepts to their biological functions. A common way for searching shared functions among genes is to incorporate the biological knowledge provided by biological ontologies [19]. The gene ontology resource is a major bioinformatics initiative that provides tools to annotate genes to their biological processes [20, 21]. We used the mouse genome informatics (MGI) term mapper to provide ontologies or biological processes of the top 10% genes or proteins [22, 23]. However, this method generated multiple functional classifications for a unique source node. Therefore, we grouped these functional ontologies by a unique ontology ID when duplicates or similar biological functions were listed. We obtained 99 genes and 117 proteins at the intersection of DKD, IR, and GEC domains and 21 unique ontology IDs that describe their biological function. The grouped genes or proteins into 21 unique biological processes are provided in Figure 6. A unique numeric label was generated for each unique ontology ID (*term label* ), and the frequency of each ontology term mapped to a source node was recorded (*term count* ).

**Figure 6.**
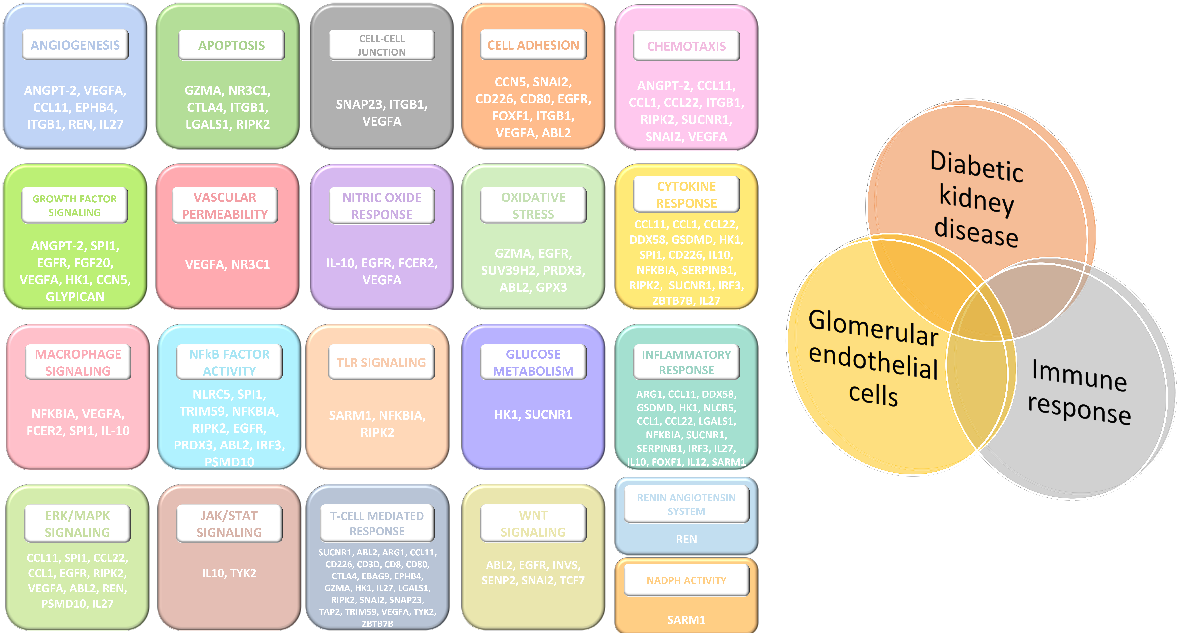
The top 10% genes and proteins that intersect in the three domains—diabetic kidney disease (DKD), immune response (IR), and glomerular endothelial cells (GEC)—are grouped into their biological processes (block headers). TLR: toll-like receptor. NADPH: nicotinamide adenine dinucleotide phosphate. NFκB: nuclear factor kappa B. WNT: wingless/integrated ERK: extracellular signal-regulated kinase. MAPK: mitogen-activated protein kinase. JAK: Janus kinase. STAT: signal transducer and activator of transcription.

Further, the biological functions associated with these identified genes or proteins are summarized. The top 10% source genes or proteins were associated with mitogen-activated protein kinase (MAPK), extracellular signal-regulated kinase (ERK1/ERK2) signaling cascade, positive regulation of receptor signaling pathway via Janus kinase/signal transducer and activator of transcription (JAK-STAT) pathway, nuclear factor-κB transcription factor activity, vascular endothelial growth factor and fibroblast growth factor signaling, Wnt signaling, and toll-like receptor signaling. These genes were found to have multiple functional roles. These genes are responsible for dysregulated glucose metabolism in cells like macrophages and immune and inflammatory responses. For instance, succinate, among the top 10% of identified source nodes, is a signaling metabolite sensed extracellularly by succinate receptor 1 (SUNCR1), and its accumulation in macrophages is known to activate a pro-inflammatory response. Similarly, Hexokinase 1 (HK1) is responsible for dysregulated glucose metabolism and is associated with cytokine response, inflammatory response, and growth factor signaling. The cross-domain analysis also identified vascular endothelial growth factor (VEGF-A) protein at the intersection of IR, GEC, and DKD domains, which is associated with mediating cell junction integrity via the VEGFR1/PLC-γ route [7]. VEGF-A is also associated with macrophage or monocyte differentiation, which suggests its role in macrophage response in pathologic conditions. Besides VEGF-A, integrin subunit beta 1 (ITGB1) is highly related to cellular junction assembly and cell adhesion. The role of some of these signaling molecules and pathways has been previously accounted for in the macrophage and GEC network model to understand the progression of early-stage DKD (Figure 1). However, performing comprehensive and integrative text mining across separate dysregulated events associated with DKD provided relevant genes and proteins that can be used to address the limitations of the existing network model.

The frequency of a biological process or function associated with a source node qualitatively describes its relative importance. Figures 7 and 8 show a heatmap of the top 10% identified source nodes and their respective biological processes specified as a unique ontology ID. The frequency of these processes is represented with a color bar on the right. These source nodes in the GEC, IR, and DKD domains play a crucial role in immune response, T-cell mediated response, cytokine response, apoptosis, and cell adhesion, among other critical biological functions (Figures 7 and 8). The biological processes like the TLR pathway, immune response, apoptotic process, calcium channel activity, and response to cytokines were relatively more prevalent processes associated with the identified source genes (Figure 7). The biological processes like T-cell mediated response, apoptotic process, cell adhesion, response to cytokines, chemotaxis, and growth factor signaling were relatively more prevalent processes associated with the identified source proteins (Figure 8). We also provided the mean HeteSim score of the identified genes (Figure S1) and proteins (Figure S2) mapped to their unique ontology ID. Genes such as SPI1, SNAP23, STMN2, and ZNF131 and proteins such as TYK2, NFKBIA, and CREG1 had the highest mean Hetesim score and are closely related to the user-specified target. CCL1, CD226, HEY2, TAP2, and TMSB10 proteins (Figure S3) frequently occurred at the intersection of DKD, GEC, and IR domains and were associated with biological processes like JAK/STAT signaling, Notch signaling, T-cell mediated response, cell migration and proliferation, response to cytokines and chemotaxis, and immune response. CD3D, CD8, LAMP2, SUV39H2, TCF7, and ZBTB7B genes (Figure S4) frequently recurred in the cross-domain analysis and were associated with T cell-mediated response, Wnt/β-catenin pathway, cell differentiation, immune response, and response to cytokines and oxidative stress.

**Figure 7.**
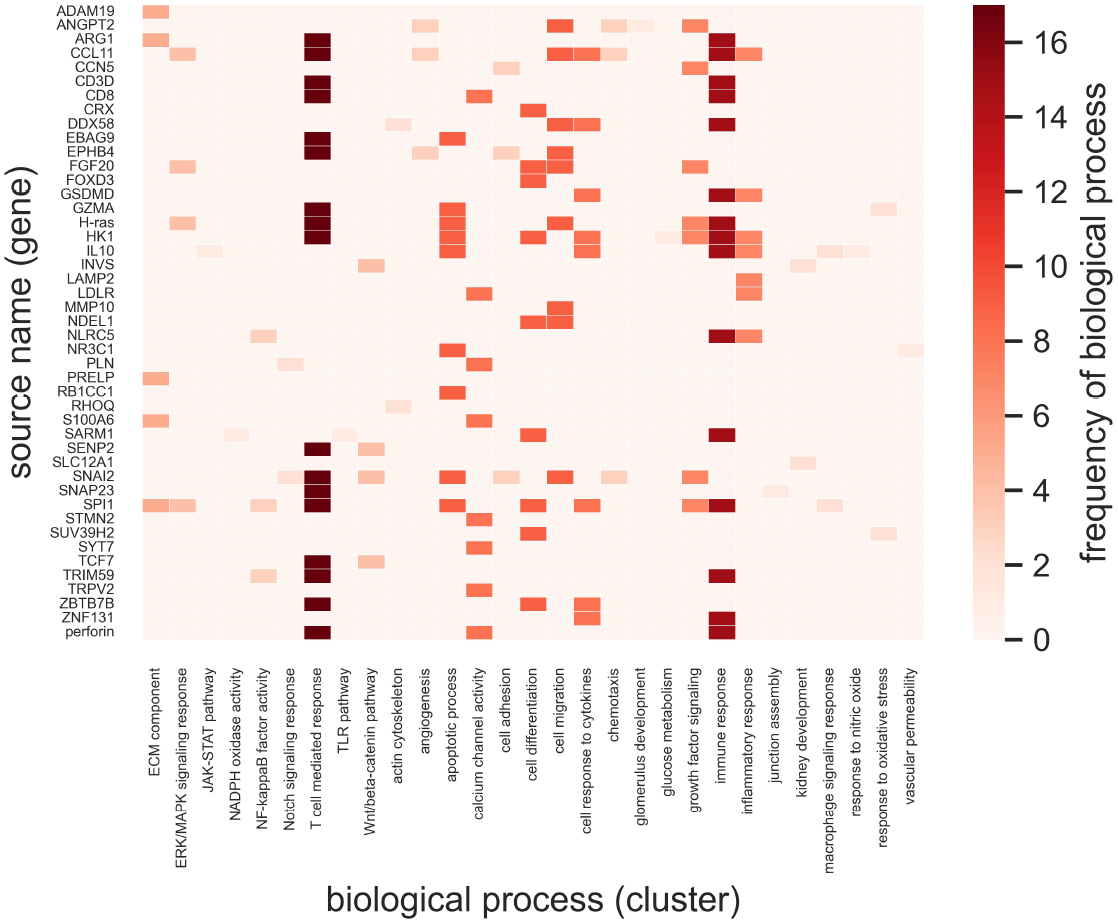
Source nodes (gene or genome) and their biological processes in the GEC, DKD, and IR domains. The frequency of each biological process is color-coded in the range of 0-16. TLR: toll-like receptor. NADPH: nicotinamide adenine dinucleotide phosphate. NFκB: nuclear factor kappa B. WNT: wingless/integrated ERK: extracellular signal-regulated kinase. MAPK: mitogen-activated protein kinase. JAK: Janus kinase. STAT: signal transducer and activator of transcription.

**Figure 8.**
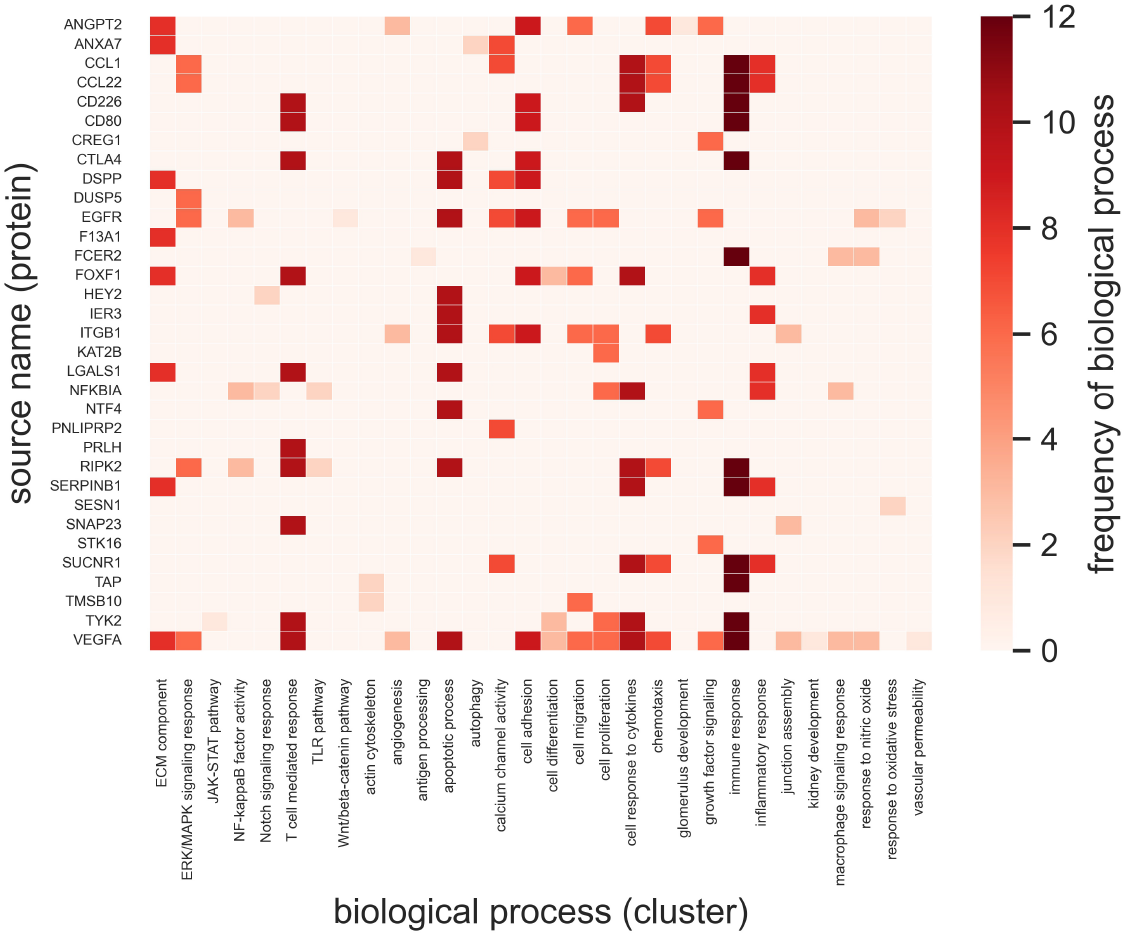
Source nodes (amino acid, peptide, or protein) and their biological processes in the GEC, DKD, and IR domains. The frequency of each process is color-coded in the range 0-12. TLR: toll-like receptor. NADPH: nicotinamide adenine dinucleotide phosphate. NFκB: nuclear factor kappa B. WNT: wingless/integrated ERK: extracellular signal-regulated kinase. MAPK: mitogen-activated protein kinase. JAK: Janus kinase. STAT: signal transducer and activator of transcription.

Cytoscape [24] was used to generate a linked protein-protein network using the identified top 10% source nodes and their mapped ontologies (Figure 9). Cytoscape is an open-source software platform for visualizing complex networks and integrating these with any attribute data [24]. We searched through the functional ontologies of the source nodes previously generated using MGI term mapper. We chose source nodes that indicated a positive or negative relationship with a signaling molecule. These relationships between source nodes and signaling molecules were selected by searching for specific words such as “positive regulation” or “negative regulation.” A protein-protein interaction (PPI) file was created to store these source nodes as inputs and signaling molecules as outputs. The interaction network was created using this PPI file (Figure 9). The interaction edges between source nodes and molecules were based on the specified positive or negative relationship using a +1 or *−*1 relation index, respectively (Figure 9). More information on generating the PPI file and format can be found in the Cytoscape user manual [25].

**Figure 9.**
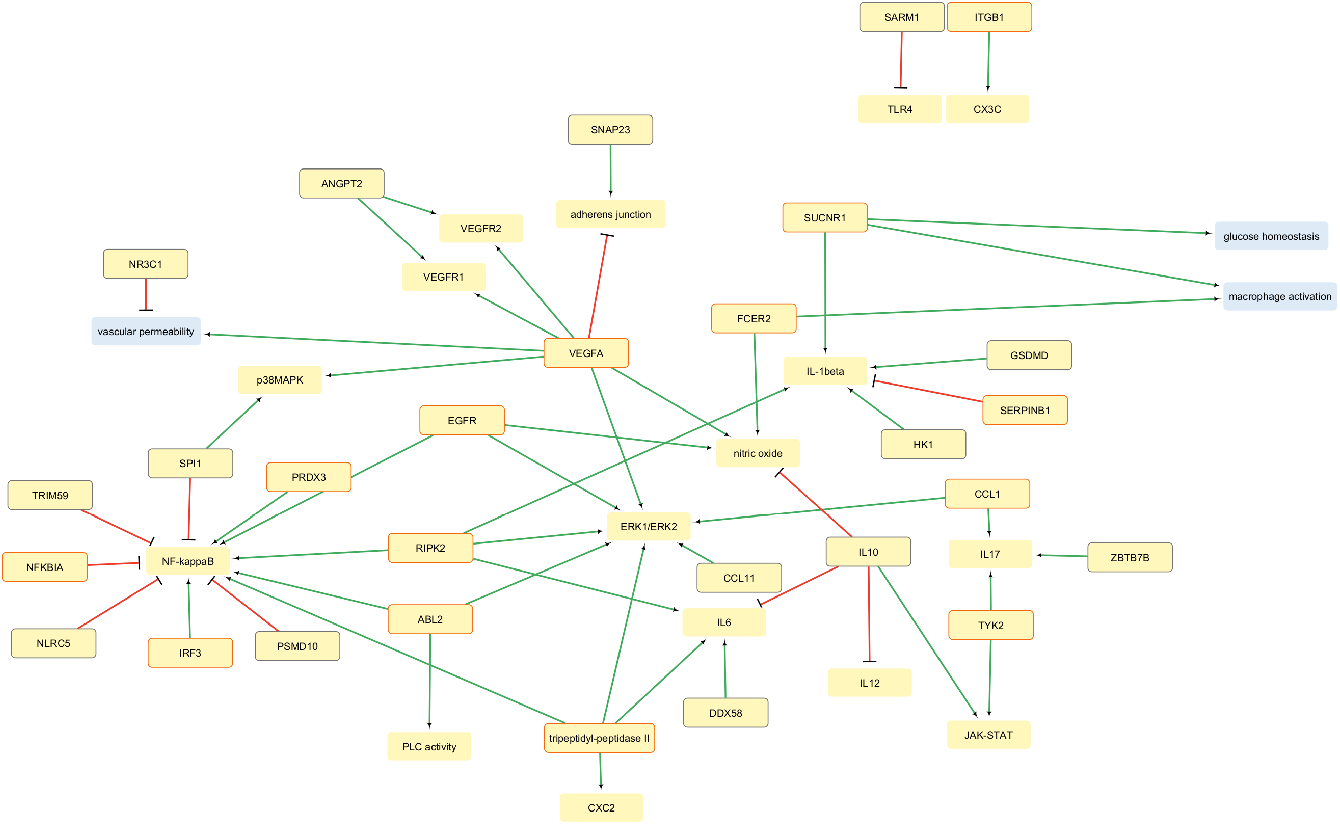
Regulation of signaling molecules by top 10% of identified source nodes in the context of diabetic kidney disease progression created in Cytoscape [28]. The green arrow is positive regulation, and the red arrow is negative regulation. Blue blocks are signaling outcomes. Yellow blocks without a border are signaling molecules. Yellow blocks with an orange border are genes, and yellow blocks (grey border) are proteins obtained from SemNet.TLR: toll-like receptor. NADPH: nicotinamide adenine dinucleotide phosphate. NF-kappaB: nuclear factor kappa B. WNT: wingless/integrated ERK: extracellular signal-regulated kinase. MAPK: mitogen-activated protein kinase. JAK: Janus kinase. STAT: signal transducer and activator of transcription.

Upon analyzing the protein-protein interaction network, we found that NF-κB, a transcription factor, is involved in numerous signaling events, including inflammatory response; it is positively regulated by PRDX3, EGFR, RIPK2, ABL2, and IRF3 genes and negatively regulated by SPI1, TRIM59, NLRC5, PSMD10 proteins, and NFKBIA gene. We found that vascular endothelial development and growth in endothelial cells heavily rely on nitric oxide, which is positively influenced by VEGF-A, EGFR, and FCER2 genes and negatively influenced by the IL-10 protein. VEGF-A growth factor positively regulates signaling molecules such as ERK1/ERK2 and p38 involved in MAPK signaling. RIPK2 and DDX58, as well as GSDMD, RIPK2, and HK1, play a positive role in regulating pro-inflammatory cytokines like IL-6 and Il-1β. Conversely, the expression of IL-6 and IL-12 cytokines is negatively regulated by the anti-inflammatory cytokine IL-10. Previous research has also demonstrated the impact of VEGF-A on vascular permeability [7, 26], which is also validated in this study.

Additionally, NR3C1 protein was found to play a role in reducing vascular permeability within endothelial cells. It was shown that the FCER2 gene was one of the identified genes involved in positive macrophage activation regulation. Moreover, the adherens junction proteins are responsible for regulating the endothelial cell-cell junction and vascular permeability in a healthy and diseased state [7, 27]. The analysis identified that adherens junction proteins are positively regulated by the SNAP23 protein and negatively regulated by the VEGF-A gene. As mentioned previously, SUCNR1 is involved in both glucose homeostasis and macrophage activation and serves as a potential link in understanding glucose-mediated macrophage cell polarization. Such a representation of source nodes using Cytoscape allowed straightforward interpretation of regulatory relationships in our data set. The comprehensive text mining analysis provided potential candidates involved with dysregulated signaling events in DKD. These identified source genes or proteins can be further studied through an interaction network-based approach and computational modeling.

## Discussion

The biomedical literature is a continuously growing repository of complex and deeply interconnected information. Despite powerful, user-friendly scientific databases, it is difficult for scientists or clinicians to extract useful information in their niche from these large and complex databases [12]. SemNet 2.0, an open-source technology applied in this study, is a literature-based discovery technique that assists scientists or clinicians by leveraging the power of literature text mining to guide their research and development efforts. In this study, novel cross-domain text mining with SemNet 2.0 identified signaling drivers and pathways that modulate glomerular endothelial dysfunction in DKD. The pairwise cross-domain provided a set of relevant genes and proteins in pathological events such as immune response, glomerular endothelial dysfunction, and diabetes, which are linked to DKD (Figure 4). SemNet 2.0 also provided a source node clustering by examining similarities between different source nodes and their specific target node in each pairwise domain (Figure 5). The source node clustering enables the user to assess potential physiological concepts such as genes or proteins with respect to the target node. It allows a better understanding of the semantic network. It helps the user to determine if a user-specified target node is sufficient to generate relevant source nodes that are not too generic.

Further, we used ontology annotators to map the top 10% of the identified genes and proteins to their biological processes. Such a functional mapping enabled the understanding of pathophysiological function, signaling pathways, and cellular events most related to disease modulation (Figures 6–8). The identified source nodes and their biological functions that had the highest association with the regulation of DKD, immune response, and GEC were selected. Further, we demonstrated the use of SemNet 2.0 analysis and prediction in generating a protein-protein interaction network through Cytoscape. Cytoscape can build network models of interaction and tools for annotating and analyzing the connections or relationships in a data set [28]. The architecture is flexible, and input data can be genes, proteins, chemicals, or enzymes [28]. CompositeView has many similarities to Cytoscape and has been successfully implemented and customized to examine SemNet or SemNet 2.0 results [29].

The study produced two valuable outcomes: the associated source nodes and the regulatory networks potentially involved in immune-mediated glomerular endothelial dysfunction in DKD.

Biomedical text mining and similarity-based clustering analyses have limitations. The clustering of these biomedical concepts or nodes based on similarity represents similarity in patterns of associations to the user-specified target node. Thus, the similarity-based association of source to target depends on the amount and quality of literature data. The implementation of additional link prediction algorithms with SemNet 2.0, as was done by McCoy and colleagues to use SemNet to predict COVID-19 drugs while the virus was new and had minimal literature [12], is one way to overcome this limitation. Regardless, a larger sample size of data reduces any bias from any lesser-quality publications. On the other hand, the user can control the loss of information when less evidence for a subject matter is available in the literature.

## 5 Conclusions

This study demonstrated that cross-domain analysis using text mining techniques provides an appropriate and efficient way to identify and prioritize relevant signaling molecules and pathways associated with DKD. The signaling molecules and pathways were identified based on the highest similarity score with the user-specified target. Many of these identified molecules were previously known to be closely related to separate pathophysiological conditions like kidney disease, diabetes, and immune response. Through our cross-domain analysis, we identified the top 10% of molecules with recurrent biological functions at the intersection of these pathophysiological domains. Including comprehensive biomedical literature in the cross-domain analysis allowed breadth in mechanistic understanding of disease progression, which is often not achievable through manual literature searches. The results from the DKD case study provide practical insights for developing protein-protein interactions and biochemical models, testing hypotheses through experiments, and advancing decision-making.

## Acknowledgments

This work was supported by National Institutes of Health grant R35GM133763 to A.N.F.V., National Science Foundation CAREER grant 2133411 to A.N.F.V., National Science Foundation CAREER grant 1944247 to C.S.M., National Institute of Health grant U19-AG056169 sub-award to C.S.M., and by the Chan Zuckerberg Initiative under grant 253558 to C.S.M.

## Supporting Information

**Figure S1.**
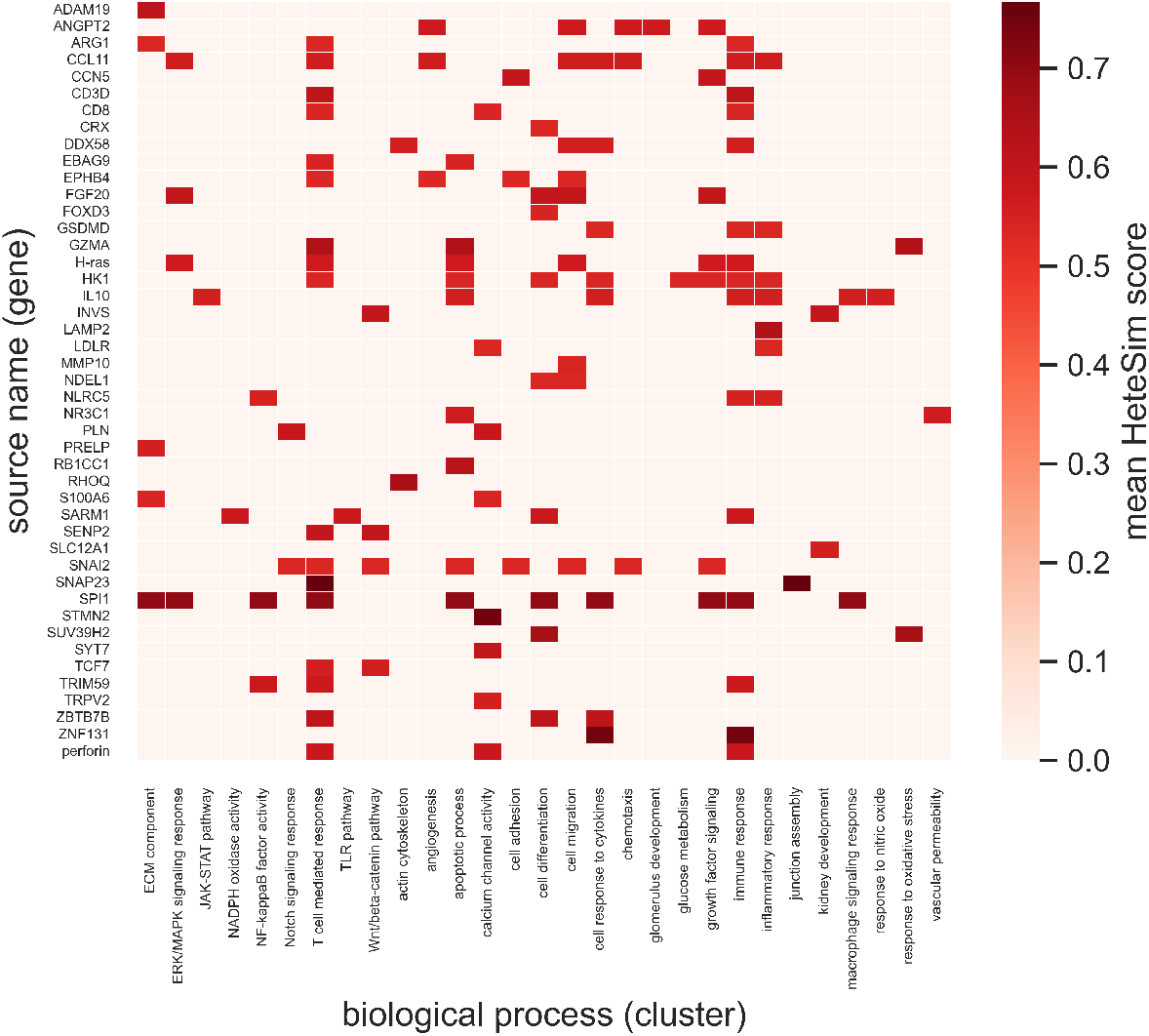
Source nodes (gene or genome) and their biological processes in the GEC, DKD, and IR domains. The mean Hetesim score of each gene from the SemNet simulation is represented by the color bar on the right. TLR: toll-like receptor. NADPH: nicotinamide adenine dinucleotide phosphate. NFκB: nuclear factor kappa B. WNT: wingless/integrated ERK: extracellular signal-regulated kinase. MAPK: mitogen-activated protein kinase. JAK: Janus kinase. STAT: signal transducer and activator of transcription.

**Figure S2.**
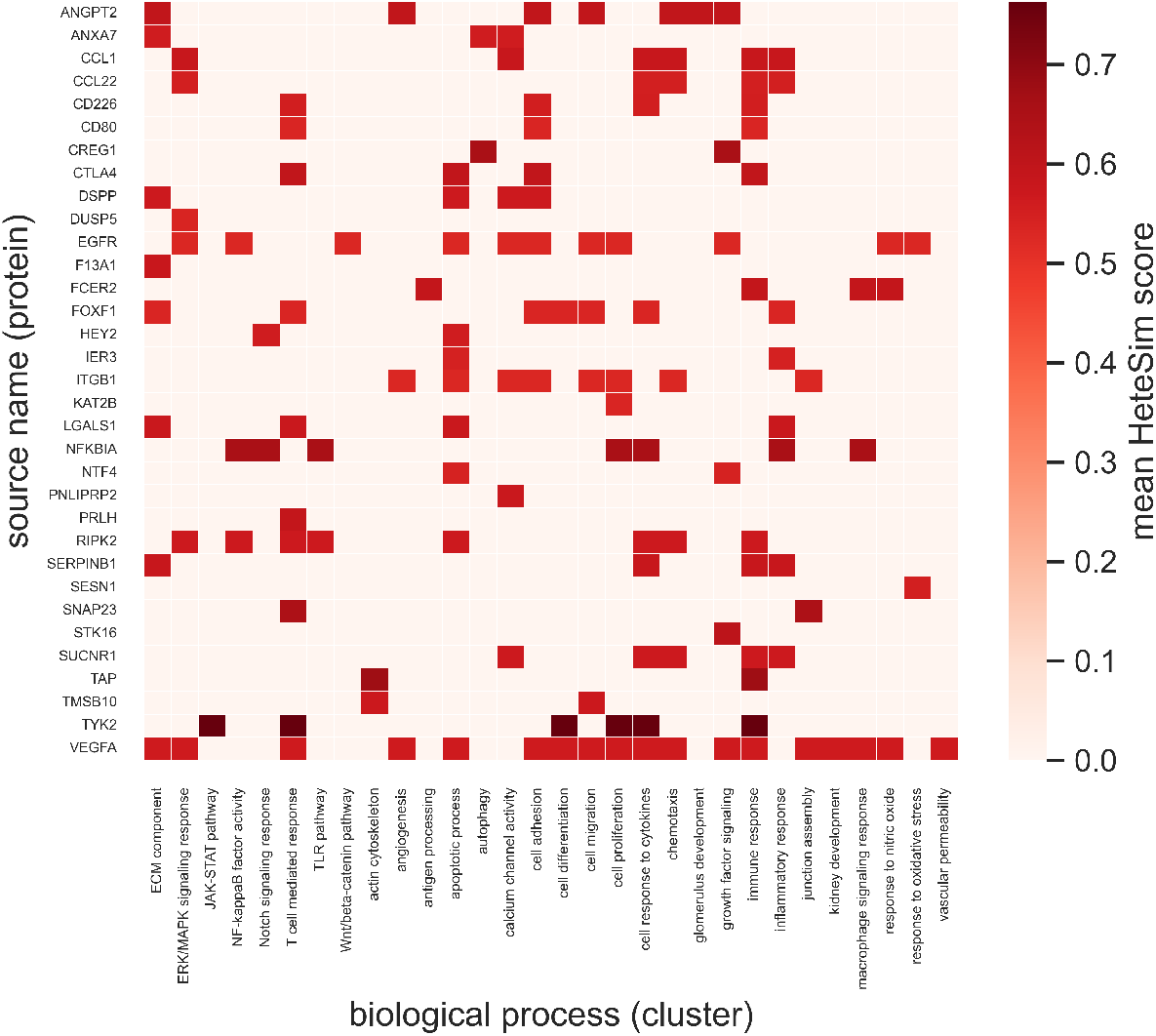
Source nodes (amino acid, peptide, or protein) and their biological processes in the GEC, DKD, and IR domains. The mean Hetesim score of each protein from the SemNet simulation is represented by the color bar on the right. TLR: toll-like receptor. NADPH: nicotinamide adenine dinucleotide phosphate. NFκB: nuclear factor kappa B. WNT: wingless/integrated ERK: extracellular signal-regulated kinase. MAPK: mitogen-activated protein kinase. JAK: Janus kinase. STAT: signal transducer and activator of transcription.

**Figure S3.**
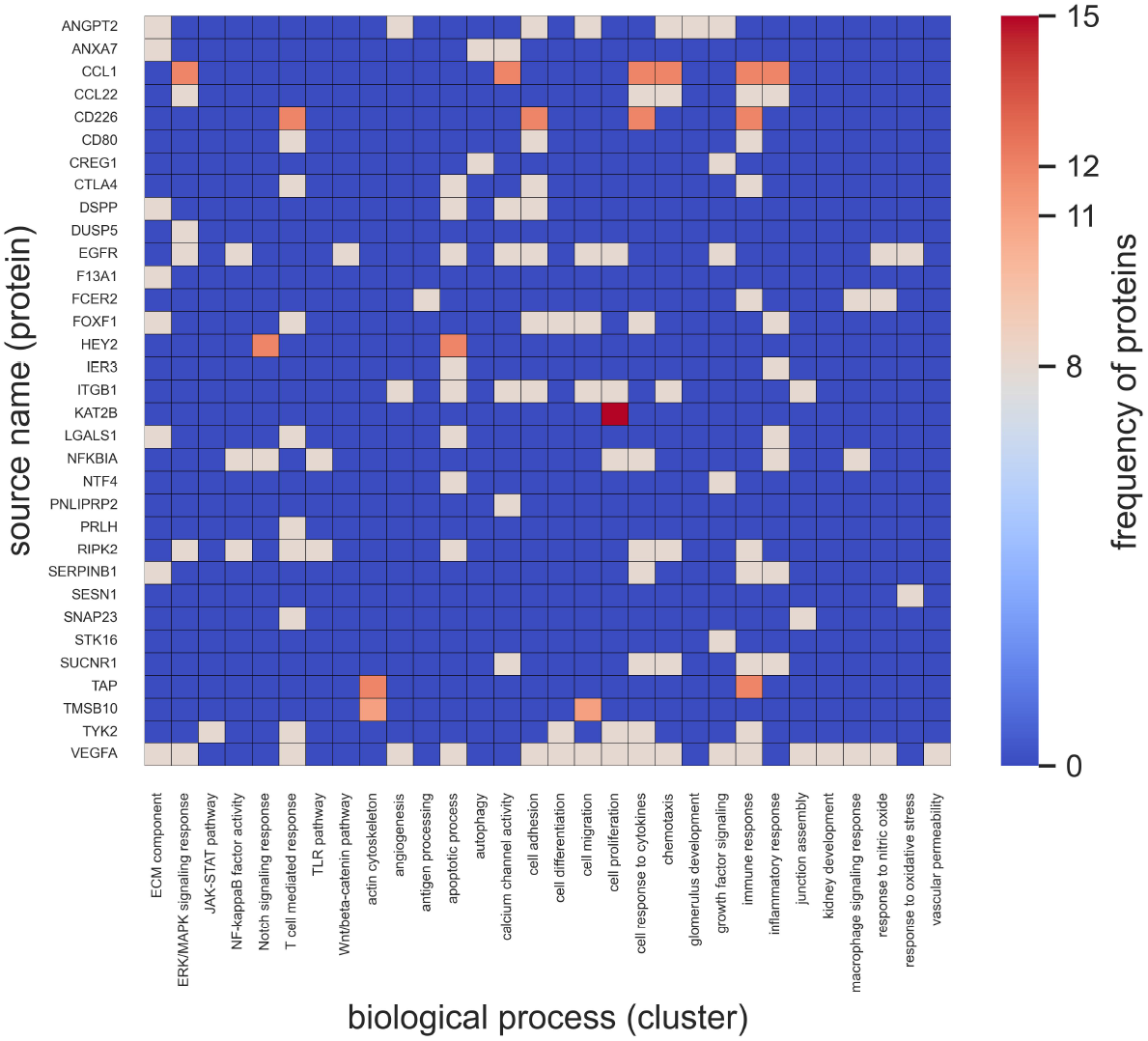
Source nodes (amino acid, peptide, or protein) and their biologicFal processes in the GEC, DKD, and IR domains. The frequency of each source protein (count) is color-coded. TLR: toll-like receptor. NADPH: nicotinamide adenine dinucleotide phosphate. NFκB: nuclear factor kappa B. WNT: wingless/integrated ERK: extracellular signal-regulated kinase. MAPK: mitogen-activated protein kinase. JAK: Janus kinase. STAT: signal transducer and activator of transcription.

**Figure S4.**
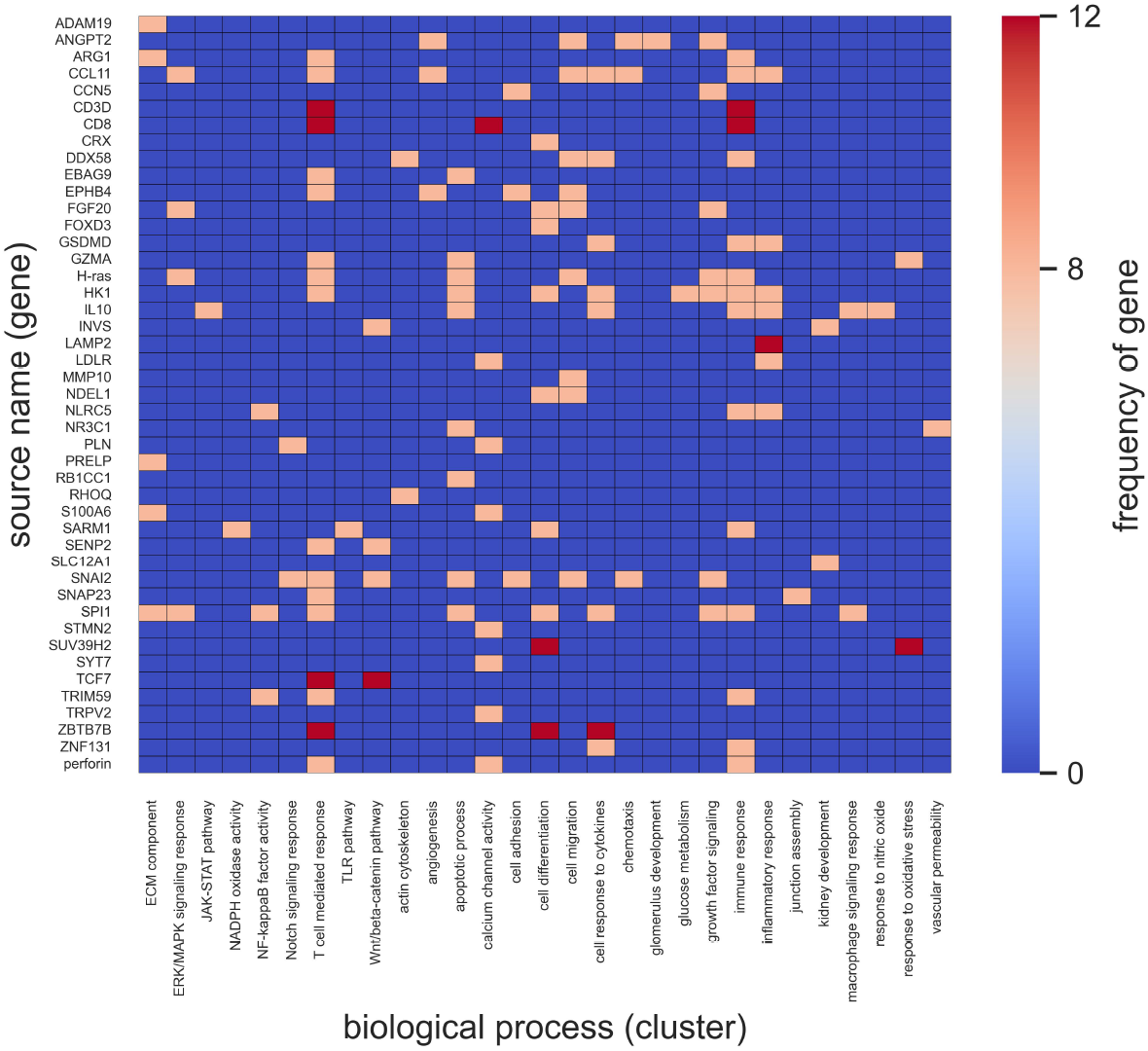
Source nodes (gene or genome) and their biological processes in the GEC, DKD, and IR domains. The frequency of each source gene (count) is color-coded. TLR: toll-like receptor. NADPH: nicotinamide adenine dinucleotide phosphate. NFκB: nuclear factor kappa B. WNT: wingless/integrated ERK: extracellular signal-regulated kinase. MAPK: mitogen-activated protein kinase. JAK: Janus kinase. STAT: signal transducer and activator of transcription.

